# AMPAR/TARP stoichiometry differentially modulates channel properties

**DOI:** 10.1101/2019.12.16.877506

**Authors:** Federico Miguez-Cabello, Nuria Sánchez-Fernández, Natalia Yefimenko, Xavier Gasull, Esther Gratacòs-Batlle, David Soto

## Abstract

AMPARs control fast synaptic communication between neurons and their function relies on auxiliary subunits, which importantly modulate channel properties. Although it has been suggested that AMPARs can bind to TARPs with variable stoichiometry, little is known about the effect that this stoichiometry exerts on certain AMPAR properties. Here we have found that AMPARs show a clear stoichiometry dependent modulation although AMPARs still need to be fully saturated with TARPs to show some typical TARP-induced characteristics (i.e. an increase in channel conductance). We also have uncovered important differences in the stoichiometric modulation between calcium-permeable and calcium-impermeable AMPARs. Moreover, in heteromeric AMPARs, TARP positioning in the complex is important to exert certain TARP-dependent features. Finally, by comparing data from recombinant receptors with endogenous AMPAR currents from cerebellar granule cells, we have determined a likely functional stoichiometry of 2 TARPs associated with GluA2 subunits in the somatic AMPARs found in this cell type.

## Introduction

Glutamate is a crucial neurotransmitter in the central nervous system (CNS) since it mediates the vast majority of the fast-excitatory synaptic transmission acting on postsynaptic ionotropic glutamate receptors. Among these, α-amino-3-hydroxy-5-methyl-4-isoxazolepropionic acid (AMPA) receptors (AMPARs) are fundamental players in the synaptic course besides their pivotal role as regulators of synaptic plasticity. AMPARs are homo or heterotetrameric structures composed of four GluA subunits (GluA1-4) (Traynelis et al. 2010) and their biophysical properties are dramatically changed depending on the subunits that conform the ion channel, with the presence or absence of the GluA2 subunit a critical determinant of channel properties. In particular, while GluA2-lacking AMPARs are permeable to both Na^+^ and Ca^2+^ ions, AMPARs containing the GluA2 subunit are impermeable to Ca^2+^. GluA2-containing AMPARs do not allow Ca^2+^ influx through the channel due to an RNA editing process that results in the replacement of an Arginine with a Glutamine residue in the cation selectivity filter region of this subunit at the so-called Q/R site (Hollmann et al. 1991; Burnashev et al. 1992). This RNA editing process is exclusive for this GluA subunit and occurs in 99% of native GluA2 subunits (Geiger et al. 1995; Kawahara et al. 2003). This change not only affects Ca^2+^ permeability but also single-channel conductance, which is decreased in GluA2-containing AMPARs mainly due to the lack of Ca^2+^ permeability (Swanson et al. 1995). This editing at the Q/R site also impairs the blocking effect of endogenous polyamines on these receptors at depolarized membrane potentials compared to GluA2-lacking AMPARs, which are strongly blocked by spermine (Bowie and Mayer, 1995; Kamboj et al. 1995; Koh et al. 1995). Thus, AMPARs are frequently classified as Ca^2+^-impermeable *vs.* Ca^2+^-permeable (CI *vs.* CP-AMPARs – or GluA2-containing *vs.* GluA2-lacking. Finally, the Q/R editing of the GluA2 subunit powerfully influences AMPAR tetramerization and strongly disfavours formation of GluA2 homotetramers (Greger et al. 2003).

Although AMPAR gating – and also trafficking – properties are determined by their subunit composition, these features are also strongly dependent on AMPAR-associated transmembrane proteins that behave as auxiliary subunits of the receptor. In the last 15 years, the number of interacting proteins that can act as modulatory partners of AMPARs has vastly increased. Stargazin and other TARPs (*Transmembrane AMPAR Regulatory Proteins*) were the first discovered proteins modulating AMPARs (Chen et al. 2000; Tomita et al. 2005). TARPs are one of the most studied AMPAR auxiliary proteins because of their indispensable role in neuronal physiology (*see (Payne 2008) for review*) and in different types of synaptic plasticity (Rouach et al. 2005; Louros et al. 2014; Sullivan et al. 2017; Louros et al. 2018). It is well-known that members of the TARP family can modify biophysical properties of AMPARs by increasing conductance, slowing down kinetics or diminishing polyamine block in CP-AMPARs (Priel et al. 2005; Soto et al. 2007). Furthermore, TARPs can modify AMPAR pharmacology such as kainate-evoked responses (Kott et al. 2007; Turetsky et al. 2005). However, much less is known about the number of TARP molecules that can be present in an AMPAR complex although AMPAR/TARP stoichiometry has been investigated by single-molecule subunit counting (Hastie et al. 2013) and electrophysiological studies that have demonstrated that AMPAR efficiency to kainate vary depending on the number of TARPs present in the receptor (Shi et al. 2009). More recently, functional studies together with high definition structural data have provided evidence for a favoured 2-TARP per AMPAR stoichiometry (Dawe et al. 2019; Herguedas et al. 2019). However, with only a small number of studies investigating AMPAR/TARP stoichiometry it is still not established whether TARPs can modulate AMPARs in a stoichiometry dependent manner.

In the present work we have determined how the number of TARP molecules per AMPAR complex modify the biophysical properties of these receptors in a stoichiometry dependent manner in both CP and CI-AMPARs. To approach this issue, we have used AMPAR/TARP fusion proteins to fix stoichiometries of 2 and 4 TARPs per AMPAR. Our results show a complex stoichiometry-dependent modulation with important differences observed between CP and CI-AMPARs. Moreover, we have attempted to reveal the endogenous functional AMPAR/TARP stoichiometry in cerebellar granule cells (CGCs) by correlating results obtained in recombinant AMPARs with recordings on CGC cultures. We have taken advantage that this cerebellar neuronal type expresses a limited variety of GluA and TARP subunits together with a lack of cornichons expression on the plasma membrane (another important AMPAR auxiliary protein) (Schwenk et al. 2009; Shi et al. 2010). We propose that 2 TARPs attached to GluA2 subunits determine functional somatic AMPAR properties in CGCs.

## Results

### TARPs induce a gradual change in CP-AMPAR kinetics

AMPAR biophysical behaviour is strongly modulated by TARPs, which differentially control AMPAR gating (Straub and Tomita, 2012; Haering et al. 2014; Jackson and Nicoll, 2011; Greger et al. 2017). The prototypical TARP γ2 slows deactivation and desensitization of AMPAR-mediated responses (Priel et al. 2005), as observed with the other members of the TARP family (Soto et al. 2009). However, the potential impact of different TARP stoichiometry on AMPAR function is vastly unknown. We wondered about the degree of the slowdown of CP-AMPAR responses in two different TARP stoichiometries to determine whether an increasing number of TARP subunits present into the AMPAR complex modulates channel properties in a gradual way or whether the presence of two TARP molecules is enough to provide AMPARs with a TARPed behaviour. We studied two fixed stoichiometries using the fusion protein GluA1:γ2, which comprises GluA1 and γ2 in the same protein product. The functional validation of this construct has been previously reported (Soto et al. 2014). We co-transfected tsA201 cells with GluA1 and/or GluA1:γ2 in such a way that GluA1 homomeric CP-AMPARs had zero, two or four TARPs per receptor. Then we extracted outside-out patches from transfected cells and we applied a 10 mM glutamate step during 100 ms with a piezoelectric controller to acquire fast AMPAR-mediated currents.

We analysed desensitization kinetics of GluA1homomeric receptors in the absence or the presence of 2 or 4 γ2 molecules. Desensitization of GluA1 in the presence of the saturating concentration of glutamate – measured as the weighted time constant (τ*_w_*) – showed a clear TARP dependence with an increase in desensitization time as the number of γ2 units in the complex increased (2.32 ± 0.16 ms, 3.77 ± 0.39 ms and 6.70 ± 0.41 ms for 0-TARPs, 2-TARPs and 4-TARPs respectively; p<0.01 for comparisons between all groups; student’s *t*-test; Figure 1A-B).

**Figure 1.**
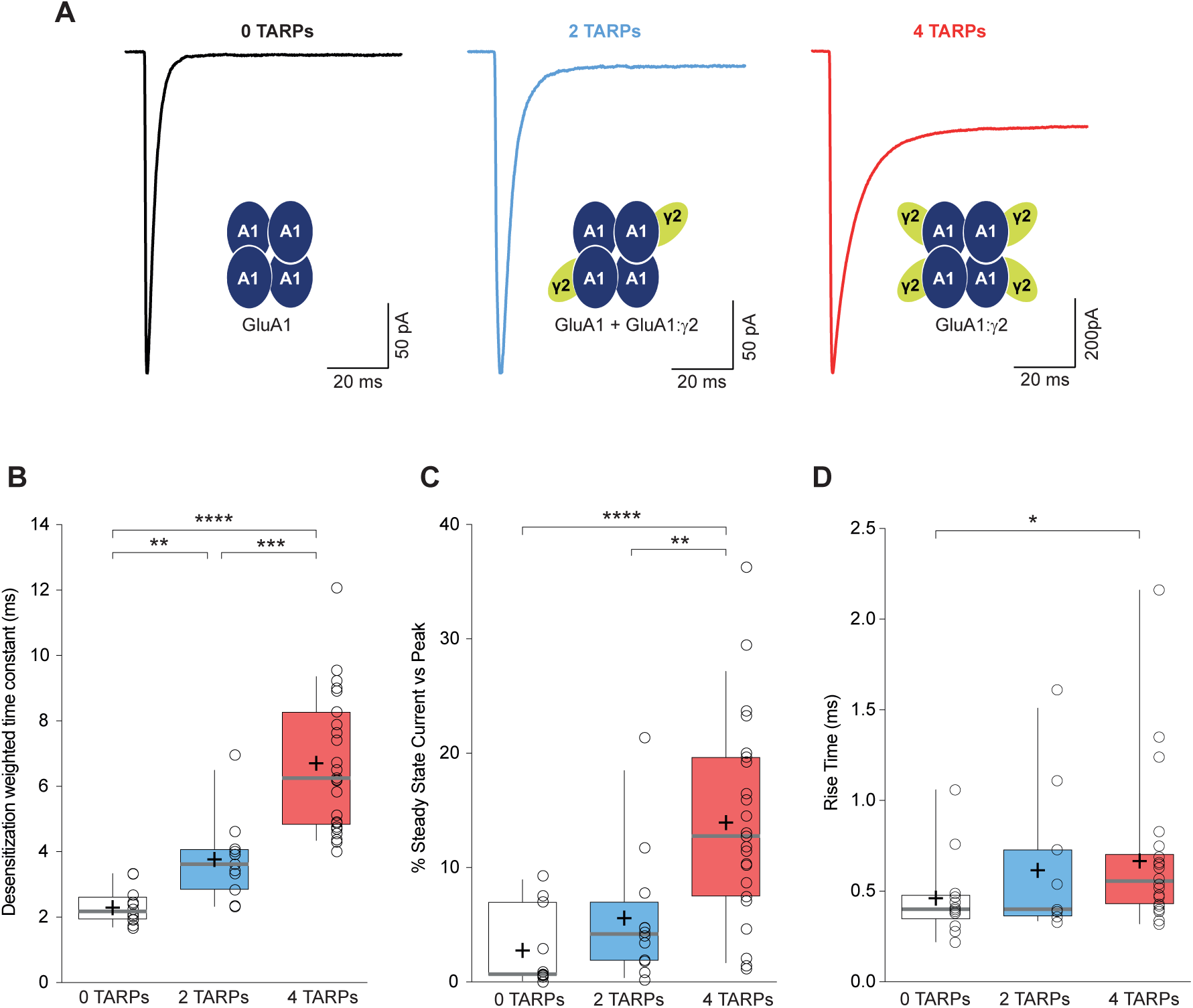
CP-AMPAR kinetics are differentially affected by AMPAR-TARP stoichiometry. A. Traces evoked at −60 mV by rapid application of 10 mM glutamate to outside-out patches from cells expressing GluA1 alone (black; average of 39 responses) or together with 2 (blue; average of 37 responses) or 4 (red; average of 67 responses) γ2 subunits. **B**. Pooled data of the weighted time constant of desensitization (τ_w, des_). Box-and-whisker plots indicate the median value (gray line), the 25–75th percentiles (box), and the 10–90th percentiles (whiskers); crosses and filled opened circles represent mean and the individual experimental values respectively (**p<0.01, ***p<0.001, ****p<0.0001). **C**. Pooled data showing the increase in the steady state current only in 4 TARPed CP-AMPARs (**p<0.01, ****p<0.0001). **D**. Rise time of glutamate-activated currents is only affected in a full TARPed CP-AMPAR (*p<0.05). The data from this figure containing statistical tests applied, exact sample number, p values and details of replicates are available in “Figure 1 - Source Data 1”.

In contrast to the well-defined step changes observed for desensitization kinetics, there was not a crystal-clear gradual change in steady state current (Figure 1C). A significant change was detected when the CP-AMPAR was fully saturated with TARPs (2.78 ± 1.04 % for 0-TARPs *vs.* 13.95 ± 1.85 % for 4-TARPs; p<0.0001; Mann Whitney *U*-test) although a gradual change cannot be discarded since a difference was observed between the 2-TARPs and 4-TARPs groups (5.58 ± 1.70 % for 2-TARPs *vs.* 13.95 ± 1.85 % for 4-TARPs; p<0.01; Mann Whitney *U*-test).

We next analysed the kinetics of the current activation (rise time) and observed that only fully TARPed GluA1 homomeric receptors presented a significant increase in the time to reach the peak current (0.46 ± 0.06 ms, 0.61 ± 0.12 ms and 0.67 ± 0.08 ms for 0-TARPs, 2-TARPs and 4-TARPs respectively; p>0.05 for all comparisons except p=0.0167 for 0-TARPs *vs.* 4-TARPs; Mann Whitney *U*-test; Figure 1D).

TARPs also speed the recovery from desensitization of AMPARs (Priel et al. 2005; Gill et al. 2012; Cais et al. 2014; Carbone and Plested, 2016) so we checked if this phenomenon was stoichiometry dependent. We applied paired pulses separated by 50 to 600 ms intervals to patches from cells expressing GluA1, GluA1+GluA1:γ2 or GluA1:γ2. Figure 2A shows typical recordings for the three conditions mentioned above. We then calculated the desensitization recovery rate and we observed a clear gradual effect, being the 2-TARPs condition half way between the slow recovery of 0-TARPs and the quicker recovery of 4-TARPs (Figure 2B). Specifically, we found time constants (τ) of 210.58 ± 1.2 ms for 0-TARPs, 194.57 ± 1.23 ms for 2-TARPs and 187.49 ± 1.65 ms for 4-TARPs (Figure 2C; n= 8, 15 and 9 respectively; p<0.01 for all comparisons; student’s *t*-test).

**Figure 2.**
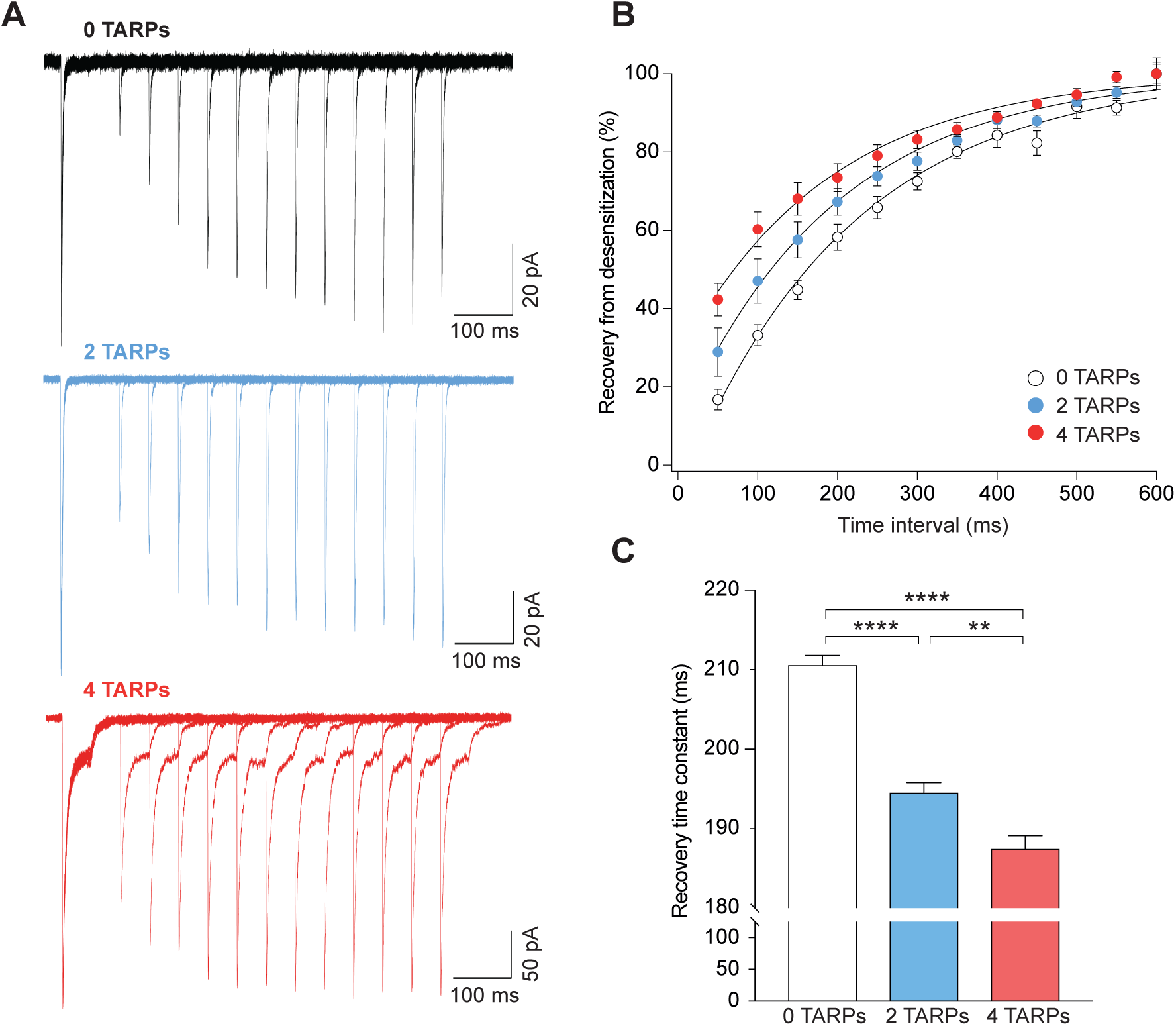
Recovery from desensitization of CP-AMPARs is gradually enhanced with increased TARP molecules. **A**. Representative traces of a two-pulse protocol with increasing time interval between pulses for CP-AMPAR without TARPs (GluA1 homomers; black), with 2 TARPs (blue) and with 4 TARPs (red). **B**. Recovery from desensitization dynamics where it can be observed the gradual diminishment in the time needed to recover as the TARP dosage increases. **C**. Recovery time constant values for the experiments showed in A and B (**p<0.01, ****p<0.0001). The data from this figure containing statistical tests applied, exact sample number, p values and details of replicates are available in “Figure 2 - Source Data 2”.

### CP-AMPAR polyamine block attenuation strongly depends on TARP dosage

An important canonical property of CP-AMPARs is the strong intracellular polyamine block of the channel especially at depolarized potentials, which translates into a characteristic inwardly rectifying current-voltage relationship (Kamboj et al. 1995). This strong block by polyamines is attenuated as a consequence of TARP modulation (Soto et al. 2007; Soto et al. 2009). Thus, we investigated if this weakening of the block followed a stepwise pattern similar to the one observed for desensitization kinetics. We observed a strong dependence on the number of TARPs associated with the CP-AMPAR in such attenuation across different membrane voltages (Figure 3A-B). The rectification index (RI; +60mV/-60mV; Figure 3C) clearly demonstrates a stoichiometry dependence of TARP-mediated attenuation of spermine block (0.056 ± 0.004 for 0-TARPs; 0.128 ± 0.011 for 2-TARPs and 0.274 ± 0.021 for 4-TARPs; p<0.0001 for comparisons between all groups; student’s *t*-test).

**Figure 3.**
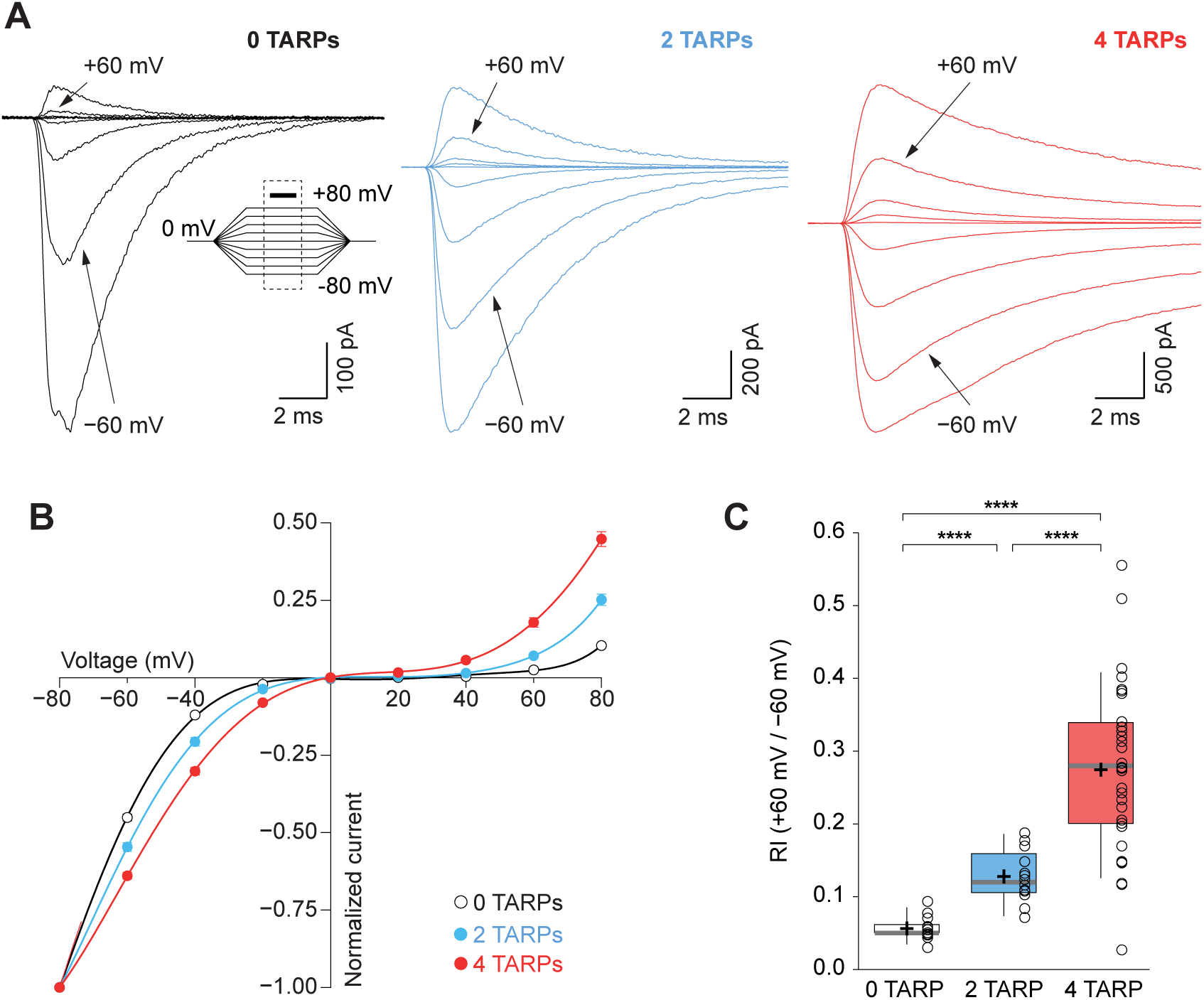
CP-AMPAR polyamine block attenuation is dependent on TARP dosage. **A**. Representative glutamate-evoked currents from outside-out patches at different membrane potentials from −80 to +80 in 20 mV increments from cells expressing CP-AMPARs, GluA1 (black), GluA1+GluA1: γ2 (blue) and GluA1: γ2 (red). Bottom: Traces at +60 mV and −60mV membrane potentials are marked. **B**. I-V relationships constructed from glutamate-evoked peak currents of patches held at different membrane potentials in different AMPAR-TARP stoichiometries. **B**. Pooled data showing an increase in the RI as the number of TARPs per CP-AMPAR increases. The RI in 2 (blue) and 4 (red, 4 TARPed) TARPs per CP-AMPAR complex is higher compared with 0 TARPs (white, TARPless) (****p<0.0001). Box-and-whiskers plots meaning as in Figure 1. The data from this figure containing statistical tests applied, exact sample number, p values and details of replicates are available in “Figure 3 - Source Data 3”.

Given the results obtained in Figures 1-3, which supports the idea of a gradual modulation of GluA1 homomeric receptors depending on the number of TARPs, we wondered whether in the studied 2-TARPed condition a mixture of two distinct populations of AMPARs – 0-TARPed and 4-TARPed – rather than a pure population of heteromeric GluA1-GluA1:γ2 existed. This would account for the intermediate phenotype observed in the recordings. Therefore, we decided to transfect tsA201 cells with GluA1(Q) together with its edited variant GluA1(R) and use the polyamine block as an indicator of the existence of an heteromeric population. GluA1(Q) homomers are strongly blocked by polyamines and since GluA1(R) homomers are strongly disfavoured, a linear response would be indicative of a GluA1(Q)-GluA1(R) heteromeric receptor. We set the RI cut point as 0.7, considering a lower value in the responses as evidence for a considerable contamination with GluA1(Q) homomers. We found that 80% of the responses showed a RI >0.7 (8 out of 10 recordings). Importantly, when we co-transfected GluA1(Q):γ2 with GluA1(R) we observed RIs above 0.7 in all recorded patches (n=9), a percentage that did not differ from the TARPless condition (p=0.4737; Fisher’s exact test). We examined the desensitization kinetics of these recordings to confirm the effect of γ2 in the complex. As detected with GluA1(Q) forms, in the GluA1(R)-containing receptors the weighted time constant (τ*_w_*) of the TARPed receptor was slowed significantly compared with the TARPless receptor (2.87 ± 0.47 for 0-TARPs *vs.* 4.32 ± 0.46 for 2-TARPs; p<0.05; student’s *t*-test) (data not shown) indicating that the results seen in the 2-TARPed conditions with GluA1(Q) were acquired most probably from a 2-TARPed population rather than two populations of TARPless and fully TARPed AMPARs.

### CP-AMPARs conductance increase by TARPs needs a fully saturated receptor

We next performed non-stationary fluctuation analysis (NSFA) to obtain single-channel conductance and peak open probability from these macroscopic responses. Figure 4A shows representative responses from TARPless, 2 TARPed and 4 TARPed AMPARs together with their corresponding NSFA (Figure 4B). We obtained conductance values for TARPless GluA1 homomeric receptors (16.58 ± 0.69 pS; Figure 4C) similar to conductance values of 16.53 pS we previously described in the laboratory (Gratacos-Batlle et al. 2014). As expected GluA1:γ2 fusion protein (4-TARPed) responses were increased ∼1.4-1.5 fold as previously described when GluA1 was co-transfected together with γ2 (Soto et al. 2014) (16.58 ± 0.69 pS for 0-TARP *vs.* 24.34 ± 1.69 pS for 4-TARP; p=0.0019; n=10 and 16 respectively; Figure 4B-C). Surprisingly, the 2-TARP condition (co-transfection of GluA1 and GluA1:γ2) did not increase single-channel conductance (16.58 ± 0.69 pS for 0-TARP *vs.* 17.03 ± 2.06 pS for 2-TARP; p=0.8373 student’s *t*-test; n=10 for both conditions), indicating that 2 TARPs were not sufficient to increase AMPAR conductance.

**Figure 4.**
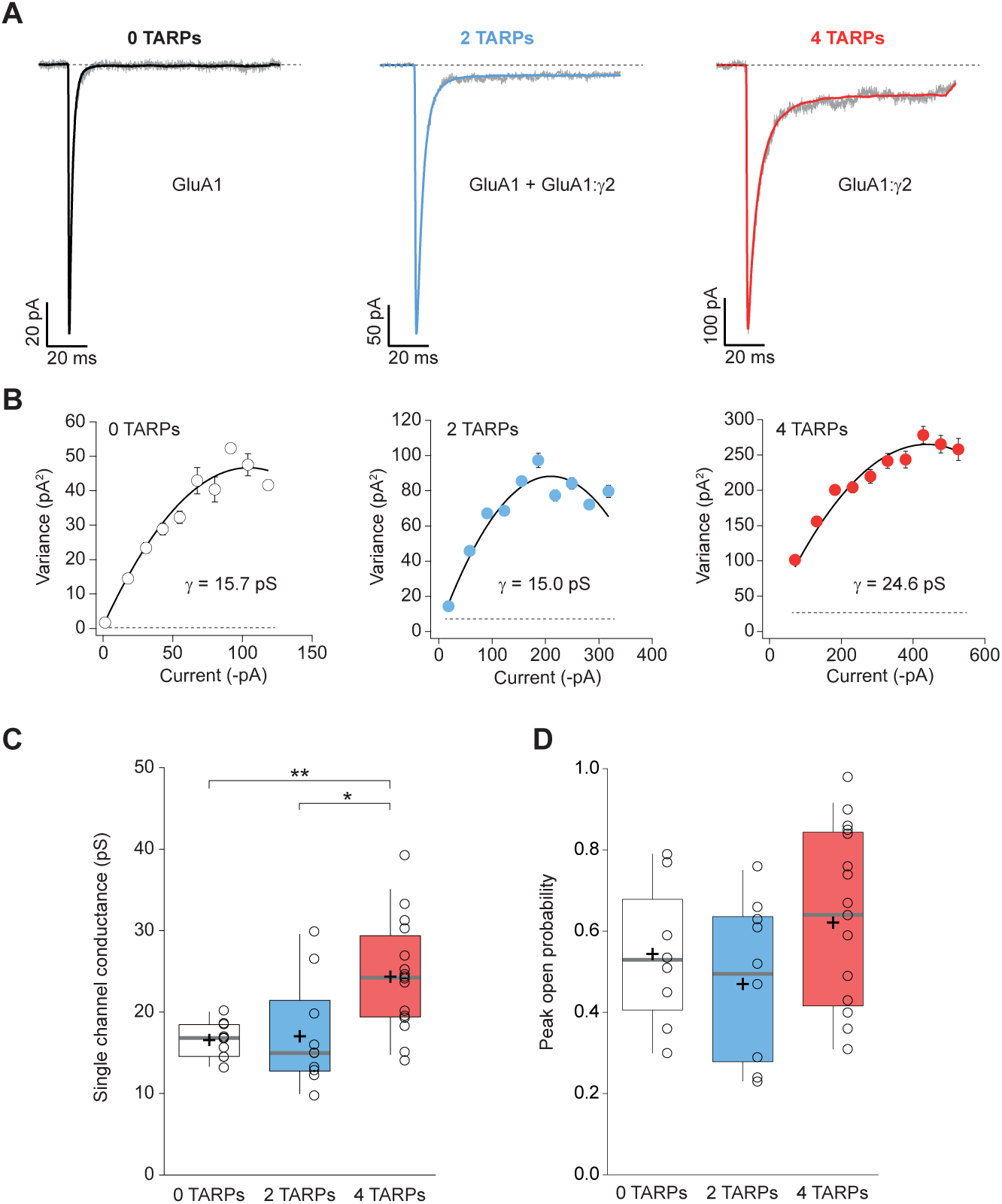
Four TARPs are required to increase CP-AMPAR channel conductance. **A**. Typical responses at a holding potential of −60 mV to rapid application of 10 mM glutamate to excised patches from cells expressing homomeric GluA1 alone (black; average of 84 responses) or together with 2 (blue; average of 91 responses) or 4 (red; average of 223 responses) γ2 subunits. A single trace is shown in gray overlaid with the mean response. **B**. Current-variance plots for the traces shown in A, the slope of which gave the weighted single-channel conductance. Broken lines show the baseline variance and error bars denote SEM. Single channel conductance values for these recordings are presented. **C**. Pooled data showing an increase of the single channel conductance only in a full-TARPed CP-AMPAR (*p<0.05, **p<0.01). **D**. Pooled data for peak open probability of CP-AMPARs. The data from this figure containing statistical tests applied, exact sample number, p values and details of replicates are available in “Figure 4 - Source Data 4”.

While it is true that TARPs have a profound effect on single channel conductance of AMPARs (Soto et al. 2007; Soto et al. 2009) their effect on the open probability is more controversial. Some studies have shown an increase of open probability in AMPARs mediated by γ2 (Suzuki et al. 2008) while others have not observed such an increase with γ2 nor with other members of the TARP I subfamily (Soto et al. 2009; Shi et al. 2009). The NSFA also allows determination of the number of channels contributing to a given response besides of the unitary conductance. Henceforth, peak open probability (P_o,peak_) can be easily deduced from the experimental mean peak current analysed. We therefore determined the P_o,peak_ of CP-AMPARs when 2 or 4 TARPs were forming part of the complex. As shown in Figure 4D, we did not find any increase of P_o,peak_ regardless of the amount of TARP in the AMPAR complex (0.54 ± 0.06 for 0-TARP *vs.* 0.47 ± 0.06 for 2-TARPs; p=0.3925 student’s *t*-test; n=9 and 10 respectively) and (0.54 ± 0.06 *vs.* 0.62 ± 0.05 for 4-TARPs; p=0.3691 student’s *t*-test; n= 9 and 17 respectively). Additionally, no change between the 2 different stoichiometries was observed (0.47 ± 0.06 for 2-TARPs *vs.* 0.62 ± 0.05 for 4-TARPs; p=0.0885 student’s *t*-test; n=10 and 17 respectively).

### Characterization of GluA4c

In order to study the effect of stoichiometry on heteromeric GluA2-containing CI-AMPARs, we focused on GluA2/GluA4, the putative AMPAR present in CGCs (Mosbacher et al. 1994). However, an alternative splicing short isoform of GluA4 – GluA4c – is highly expressed in CGCs (Gallo et al. 1992; Kawahara et al. 2004) and we thus considered that GluA2/GluA4c would be a better readout of CGC AMPARs. Consequently, the examination of this specific combination would permit us to study CI-AMPARs and compare later on the results from expression systems with data extracted from CGCs.

We first explored the behaviour of homomeric GluA4c(i) (referred hereafter as GluA4c) receptors given the relative scarcity of data with regard to GluA4c in the literature. When both forms of GluA4 were compared we observed that GluA4c homomeric AMPARs were functionally identical to GluA4 homomers. No differences were spotted in the single channel conductance or peak open probability (Figure S1A-B) measured by means of NSFA. Likewise, desensitization kinetics did not differ between both isoforms (Figure S1C). Finally, both homomeric receptors presented the same degree of intracellular block by spermine (Figure S1D-E).

### CI-AMPAR single channel conductance is differentially affected depending on TARP location within AMPAR complex

Once GluA4c properties were validated, we created a GluA4c:γ2 fusion protein to study AMPAR/TARP stoichiometry in CI-AMPARs. Unlike homomeric GluA1 receptors, in a heteromeric receptor such as GluA2/GluA4c, a 2-TARPed configuration might be achieved by locating the TARPs either in the GluA2 or in GluA4c subunit. Since this might be relevant, we co-transfected GluA2, GluA4c, GluA2:γ2 or GluA4c:γ2 to obtain 0, 2-TARPed in GluA4c, 2-TARPed in GluA2 or a fully TARPed heteromeric AMPARs. We studied responses in out-side out patches where GluA2 presence into the AMPAR was evaluated with the linearity of the responses (Figure 5A).

**Figure 5.**
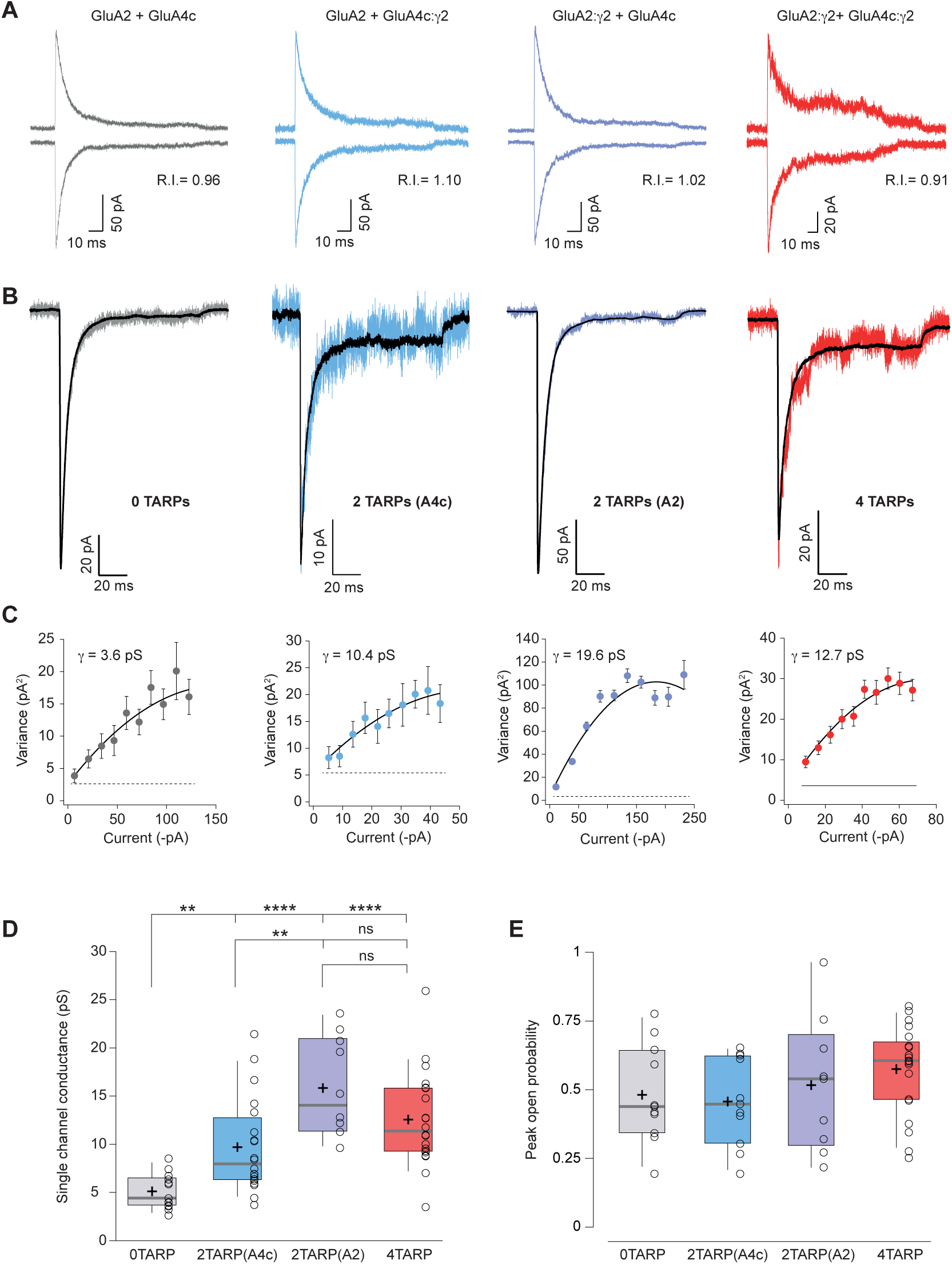
Single channel conductance of CI-AMPARs is modulated differently by TARPs depending on their location within the complex. A. Evoked currents by rapid application of 10 mM glutamate from membrane patches at +60 mV (upward traces) and −60 mV (downward traces) with their corresponding RI. The experimental conditions are designated as grey for 0 TARPs per AMPAR, blue for 2 TARPs per AMPAR with γ2 linked to GluA4c, purple for 2 TARPs per AMPAR with γ2 linked to GluA2 and red for 4 TARPs per AMPAR. **B.** Average traces of current responses evoked at −60 mV used for NSNA shown in black overlaid with a representative single response. Insets shown the studied combination. **C**. Current-variance plots for the recordings shown in B, with the weighted single-channel conductance for the single recordings. **D**. Pooled data showing a distinct degree in single channel conductance increase when γ2 is present into the AMPAR complex (*p<0.05, **p<0.01, ***p<0.001, ****p<0.0001). **E**. Pooled data for peak open probability of CI-AMPARs, where no effect of TARP stoichiometry was evident (p>0.05). The data from this figure containing statistical tests applied, exact sample number, p values and details of replicates are available in “Figure 5 - Source Data 5”.

NSFA performed on glutamate-evoked responses onto patches from tsA201 cells transfected with different combinations (Figure 5B-C) depicted a remarkable effect not observed with CP-AMPARs. Although any number of TARPs into the CI-AMPAR was sufficient to increase single channel conductance as reported previously (Jackson et al. 2011) (p<0.001 for all comparisons between 0-TARPs and the other TARPed groups; Figure 5C-D), the TARP location within the complex was important in determining the extent of conductance increase. Specifically, the TARP γ2 intensified its effect on single channel conductance when it was attached to GluA2 subunit compared with GluA4c (15.85 ± 1.62 pS *vs.* 9.72 ± 1.09 pS for γ2 attached to GluA2 or GluA4c respectively; p<0.01; student’s *t*-test; n=10 and 20 patches), which represents a 309% and an 189% of conductance increase respect to GluA2/GluA4c (5.13 ± 0.50 pS). Finally, peak open probability deducted from the same NSFA did not seem to be affected regardless of the number of TARPs or their location on GluA2-containing heteromers (Figure 5E).

### TARP γ2 linked to GluA4c slows down desensitization kinetics of CI-AMPARs

The striking observation about a differential modulation of γ2 depending on its particular position into the receptor prompted us to investigate whether other properties of CI-AMPARs were also affected differentially when γ2 was linked to one or another subunit. Figure 6A-B shows that desensitization kinetics of GluA2/GluA4c heteromeric combination (0T) were slowed down only when γ2 was linked to GluA4c subunit irrespective of a 2- or 4-TARP stoichiometry. Indeed, the kinetics of GluA2/GluA4c:γ2 – 2T(A4c) – were not different from GluA2:γ2/GluA4c:γ2 – 4T – (7.23 ± 0.43 ms for 2T(A4c) *vs.* 8.43 ± 0.61 ms for 4T; p>0.05; Mann Whitney *U*-test; n=21 and 20 respectively; Figure 6C) while both combinations were slower than 0T or 2T(A2) (p<0.05; Figure 6C). Finally, γ2 linked to GluA2 subunits did not seem to affect GluA2/GluA4c desensitization (4.76± 0.28 ms for 0T *vs.* 5.42 ± 0.40 ms for 2T(A2); p>0.05; student’s t-test; n=14 and 11, respectively). Thus, kinetic behaviour in CI-AMPARs is apparently not changed by γ2 unless the TARP is attached to GluA4c subunit. Regarding the activation time to reach the peak current (rise time), we did not notice any variation amongst the different combinations tested (Figure 6D).

**Figure 6.**
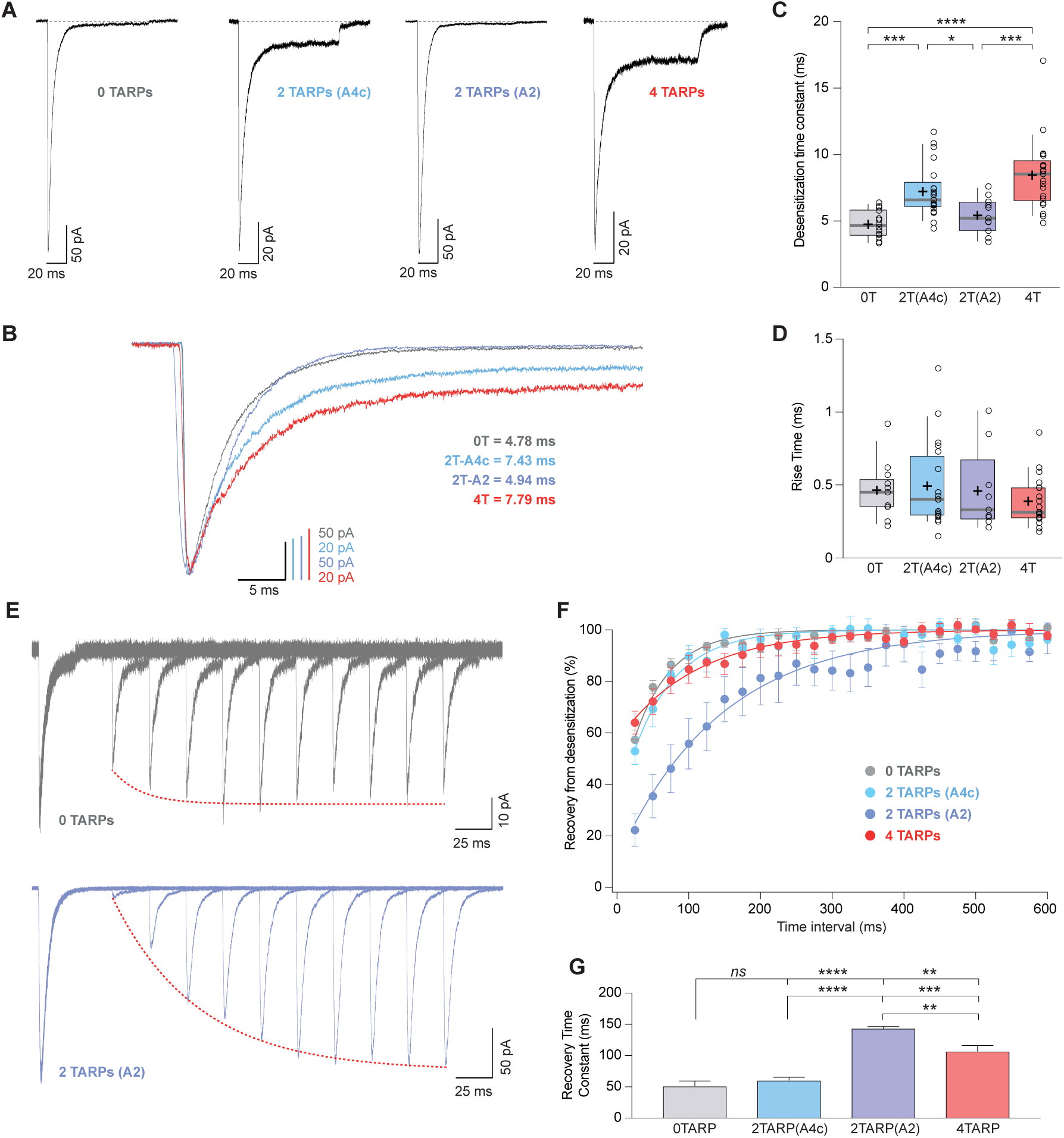
CI-AMPAR kinetics differ upon γ2 attachment to GluA4c or GluA2 subunit. **A**. Representative traces of currents at −60 mV from cells expressing CI-AMPARs without or with TARP γ2 linked to GluA subunits. Under the traces a scheme of the subunits forming the receptors with γ2 associated to different AMPAR subunits is shown. **B**. Peak-scaled normalization from traces shown in A for a better comparison of desensitization kinetics. **C**. Weighted time constant of desensitization (τ*_w,des_*) were is clear that desensitization is slowed only when γ2 is linked to GluA4c subunit (*p<0.05, ***p<0.001, ****p<0.0001).. **D**. Rise time of the current activation is not changed by the AMPAR-TARP stoichiometry. **E.** Representative traces monitoring recovery from desensitization for CI-AMPAR in cells expressing 0 TARPs or 2 TARPs linked to GluA2 subunit where it is manifest the difference between the two conditions. Dashed red line: exponential fitting for the desensitization recovery. **F**. Recovery of desensitization kinetics showing a relatively slow recovery only in 2-TARPed (located in GluA2) CI-AMPARs. **G**. Recovery time constant values for the experiments showed in E and F (**p<0.01, ***p<0.001, ****p<0.0001). The data from this figure containing statistical tests applied, exact sample number, p values and details of replicates are available in “Figure 6 - Source Data 6”.

### TARP γ2 hinders recovery from desensitization of CI-AMPARs by acting on specific subunits

We also checked how recovery from desensitization of CI-AMPARs was affected by distinct AMPAR:TARP stoichiometries. While γ2 enhanced recovery from desensitization from CP-AMPARs in a stoichiometry dependent manner, this was not the case for CI-AMPARs where faster recoveries were not observed regardless of any amount of TARP present into the AMPAR. Interestingly the opposite effect (slowest recovery from desensitization) was observed for the 2T(A2) receptor (Figure 6E-G).

### CI-AMPAR pharmacology is altered depending on their TARP stoichiometry

A previous study revealed that AMPAR pharmacology is dependent on the number of TARPs, as the efficiency of the partial agonist kainate is enhanced as the number of TARPs increase into the AMPAR complex (Shi et al. 2009). We wondered whether perampanel, a non-competitive inhibitor of AMPARs, would vary its blocking effect on AMPARs depending on the number of TARPs present. To address this question, we performed recordings where we rapidly applied perampanel in a set of experiments where whole-cell currents were activated in transfected tsA201 held at −60mV with 100 μM AMPA plus 50 μM cyclothiazide to avoid desensitization (Figure S2A). The outcome of those experiments did show the same pattern as the one found for desensitization: TARP γ2 modified the percentage of block only when attached to GluA4c subunit. CI-AMPARs with 2 TARPs at the GluA2 subunit displayed a similar block as TARPless CI-AMPARs (47.33 ± 5.95 % for 0T *vs.* 46.13 ± 5.47 % for 2T(A2); p>0.05; student’s t-test; n= 6 and 5, respectively; Figure S2C). However, the block by perampanel in a 2-TARPed at GluA4c (70.65 ± 8.06; n=7) and in a 4-TARPed CI-AMPARs (71.64 ± 6.76; n=6) was higher than for TARPless receptors (p<0.05 for 0T *vs.* 2T(A4c) and 0T *vs.* 4T; student’s t-tests).

### Somatic AMPARs from cerebellar granule cells display features of 2-TARPed AMPARs

CGCs have a high expression of GluA2 and GluA4c AMPAR subunits. Besides, these neurons only express two members of the TARPs (γ2 and γ7) (Fukaya et al. 2005) and no other auxiliary subunits such as cornichons have been described to be functionally present. While γ2 has been proved to be essential in AMPAR signalling in this cell type (Chen et al. 2000), the role of γ7 does not seem to be important to determine CI-AMPAR expression in granule cells (Studniarczyk et al. 2013). This converts CGCs into a well-defined system to study AMPARs. Thus, to determine whether CI-AMPARs in CGCs showed properties indicative of a given TARP stoichiometry, we firstly extracted somatic patches from 6-8 days in vitro CGC cultures (Figure 7A) and applied the selective agonist AMPA at 100 μM. We obtained non-rectifying responses such as the one shown in Figure 7B and performed NSFA (Figure 7C-D) to extract single channel conductance and peak open probability values from the recorded responses (Figures 7E-F; orange box) and we also calculated desensitization values from those same patches (Figure 7G; orange box). We next compared those results with the ones obtained in expression systems shown previously in figures 5 and 6. Figures 7E to G show, for each analysed parameter, the CGCs results (orange box) together with the different combinations analysed in which dull colours represent the comparable combinations (with no significant differences) and in loose grey the improbable combination responsible for currents found in CGCs (p<0.05). Taken together, it was clear that values obtained from CGCs were closely equivalent to values from recombinant receptors only for a 2-TARPed CI-AMPAR with the two TARP molecules linked to the GluA2 subunit.

**Figure 7.**
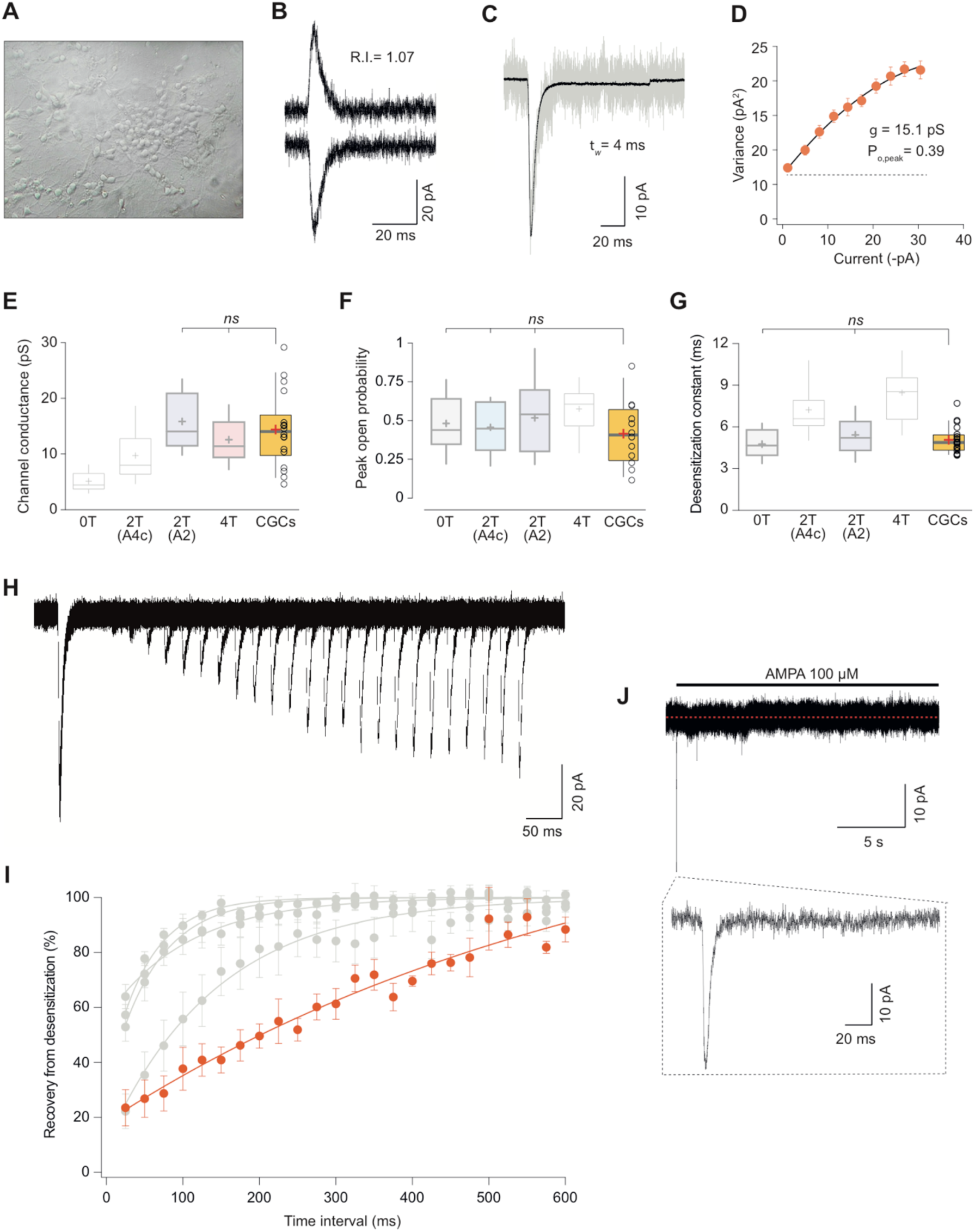
Somatic currents from CGCs exhibit properties of GluA2:γ2 + GluA4c CI-AMPARs. **A**. CGCs in culture after 7 days in vitro. **B**. Traces at +60mV and −60mV evoked with 100 μM AMPA from a CGC somatic patch showing the typical lineal response of a CI-AMPAR. **C**. Representative response of current evoked at −60 mV by rapid application of 100 μM AMPA to somatic patches from CGCs. Grey: representative single response; black: average of 275 stable responses. **D**. NSFA from the recording in C. **E**. Data showing comparison of single channel conductance values obtained in CGCs (orange box) with recordings from transfected cell lines. The conductance values obtained resembled (without significant difference) to the ones seen with 2T (A2) or 4T conditions (marked in bold grey). **F**. Comparison of peak open probability values from CGCs were only significantly different from recordings in cell lines with 4 TARPs. **G**. Compared data of desensitization time constant (ms) from CGCs with recordings in tsA201 cells. The results are no significantly different from conditions with 0T or 2T(A2). **H**. Representative trace from two-pulse protocol monitoring recovery from desensitization for CGCs somatic patches to 100 μM AMPA application. **I**. Recovery from desensitization kinetics of CGS somatic AMPARs. **J**. Representative response to a 100 μM AMPA application for 20 seconds in a somatic patch from CGCs to test for the presence of γ7. No re-sensitization of the receptors is observed in the trace. Inset: magnification of 200 ms showing the initial fast desensitizing response. The data from this figure containing statistical tests applied, exact sample number, p values and details of replicates are available in “Figure 7 - Source Data 7”.

We next studied the recovery from desensitization in CGCs and compared the results with the different CI-AMPAR combinations analysed in this study. Surprisingly, we consistently recorded currents with an exceptionally slow recovery that were different from any CI-AMPAR studied (Figure 7I). Given that no other TARP seem to be present on CGCs except γ7, we asked if this auxiliary subunit was responsible for the extraordinary slow recovery seen on AMPARs from CGCs. We examined the presence/absence of γ7 by applying a 20 second jump of AMPA to CGC somatic patches. This type II TARP has been demonstrated to confer a remarkable feature to either homomeric and heteromeric GluA2-containing AMPARs consisting in resensitization of the current a few seconds after desensitization (Kato et al. 2007; Kato et al. 2010). We did not appreciate any change in desensitized current in the presence of the agonist for the whole recording period (Figure 7J) ruling out any contribution of γ7 to CI-AMPAR currents from somatic CGCs. We also wondered about the possibility that the use of the partial agonist (AMPA) to elicit the currents might slow down the recovery from desensitization. However, recordings from cell lines studying GluA2:γ2+GluA4c did not vary when we used AMPA as agonist (data not shown) ruling out this possibility. Considering the reported absence of other members of the TARP family, the functional absence of CNIH2 (Shi et al. 2010) and the really low expression of other AMPAR auxiliary proteins as GSG1L and CKAMP44 (shisa9) (Zeisel et al. 2018) in CGCs, our results suggest that another “unknown player” might be present into the complex, which would be responsible for the extremely low recovery from desensitization. Regardless, our results seem to indicate that CI-AMPARs from somatic CGCs are very possibly modulated by two γ2 molecules attached to the GluA2 subunits.

## Discussion

One of the first evidences of a functionally variable AMPAR-TARP stoichiometry came from the observation that mEPSCs were differentially altered depending on the TARP expression levels (Milstein et al. 2007). Shortly after, by using fusion constructs similar to the ones used in our study it was seen that the pharmacology – the kainate efficiency – of recombinant AMPARs was stoichiometry-dependent (Shi et al. 2009). This same work suggested that AMPARs from hippocampal pyramidal and dentate gyrus granule neurons were 4 and 2-TARPed, respectively although a 2-TARPed complex was shown to drive AMPAR function at the main AMPAR combination found in hippocampal pyramidal neurons (Herguedas et al. 2019). Finally, recently published work has provided evidence for the presence of different stoichiometries in cerebellar cells – 2 and 4 TARPed AMPARs in stellate and Purkinje cells respectively (Dawe et al. 2019). We have expanded these previous findings by carefully dissecting the effect of different stoichiometries on AMPAR basic properties. We demonstrate a sophisticated modulation of TARPs either in CP- and CI-AMPARs and we propose that 2 TARPs acting on the gating-controlling BD “pore distal” subunit GluA2 modulates somatic AMPARs in CGCs in line with recent reports on AMPARs in other neuronal types (Herguedas et al. 2019).

### Gradual *vs*. all-or-nothing modulation of AMPARs by TARPs

We have found a clear gradual stoichiometry dependence for some of the parameters modulated by TARPs. In example, desensitization kinetics, attenuation of polyamine block by TARPs or recovery from desensitization of GluA1 homomers are differentially altered when 2 or 4 TARPs are part of the AMPAR complex. However, a more intricate modulation appears to be present since a full TARPed GluA1 receptor is necessary to vary other CP-AMPAR intrinsic properties such as single channel conductance or rise time. On the other hand, other characteristics are not altered by the number of TARPs acting on AMPARs (open probability). Thus, a multifaceted scenario arises regarding how TARPs alter AMPAR behaviour. It might be interesting in the future to test whether other auxiliary subunits from the TARP family modulate the AMPAR features studied here, especially single channel conductance since 4 copies of γ2 appear to be required to induce the typical increase of TARPs, which is well described in the literature (Soto et al. 2007; Shi et al. 2009; Soto et al. 2009; Suzuki et al. 2008; Tomita et al. 2005) and given that type Ib TARPs (γ4 and γ8) seem to associate with AMPARs in a 2-TARPed dependent manner (Hastie et al. 2013; Herguedas et al. 2019) although they clearly produce an increase in AMPAR channel conductance (Shi et al. 2010; Suzuki et al. 2008; Soto et al. 2009).

In our hands, AMPAR rise time of the current appears to be slightly affected only with the receptor saturated with four γ2 molecules. For this same type of homomeric CP-AMPAR it has been shown in HEK cells that only TARPs γ4 and γ8 are able to increase the period of time to the peak response (Milstein et al. 2007). Since these contradictory results have been obtained in the same type of expression system, the disparity in the outcome might be explained by the fact that in Milstein et al. TARPs and AMPAR subunits were transfected into the cells using two separated plasmids thus raising the possibility that AMPARs studied in the above-mentioned work were not fully saturated with γ2. However, it is interesting that in Milstein’s work, γ4 increased the rise time despite the fact that the most prevalent and favoured conformation when γ4 is the auxiliary subunit modulating GluA1 is 2-TARPed (Hastie et al. 2013), showing again the huge difference in the effects that distinct TARPs have on AMPARs. Previous mEPSCs recordings from CGCs, which are mediated by CI-AMPARs have revealed no differences in the rise time of these quantal synaptic currents when the “dose” of TARP γ2 was altered into the receptor by using homozygous or heterozygous stargazer mice (Milstein et al. 2007). In line with these results, our data of rise time of GluA2/GluA4c seems to indicate that AMPAR current activation is independent of the number of TARPs modulating the receptor in CI-AMPARs.

### GluA2 preference of TARP γ2 in native AMPARs

Our data from CGCs suggest that, in a GluA2-containing heteromeric AMPAR, γ2 preferentially binds to GluA2 subunits to modulate ion channel properties. A very recent publication by Greger’s lab has shown that in GluA1/GluA2 heteromers, the main AMPAR combination present in the hippocampus (Tomita et al. 2003; Schwenk et al. 2014), TARP γ8 binds to GluA2 subunits (Herguedas et al. 2019). It is possible that the 2-TARPed AMPAR arrangement found in this cell type might be determined by the presence of CNIH2 in the AMPARs (Gill et al. 2011; Kato et al. 2010). However, in granule cells from the cerebellum CNIH2 functional expression at plasma membrane is absent (Shi et al. 2010) and yet a 2-TARPed stoichiometry seems to be functionally present, indicating also a preferential 2-TARPed disposition at GluA2 in heteromeric GluA2-containing AMPARs. Moreover, structural studies done with γ2 suggest that TARPs preferentially bind to BD “pore distal” subunits (Twomey et al. 2016), a position preferentially occupied by GluA2 subunits (He et al. 2016; Herguedas et al. 2019) (but see (Herguedas et al. 2016)). Thus, our results strengthen the idea of a preferential arrangement of TARPs into an heteromeric complex.

### Functional AMPAR/TARP stoichiometry in granule cells from the cerebellum

Our experiments indicate that 2-TARPed AMPARs are responsible for somatic responses in CGCs. In fact, the overexpression of TARP γ2 in this cell type increased kainate affinity (Milstein et al. 2007), indicating that CGCs are not totally saturated by TARPs since the efficiency to kainate is strongly dependent on the number of TARPs in the complex (Shi et al. 2009). When comparing the data from GluA2/GluA4 heteromeric receptors using fusion proteins with those form CGCs it is evident that neither zero TARPs (low conductance) nor four TARPs (slow desensitization kinetics) are modulating somatic AMPARs in CGCs. Importantly, the findings in CGCs closely recapitulated those on expression systems only when 2 TARPs were attached to GluA2 subunit. It has been suggested that TARP subtypes might have different binding sites in the AMPAR complex (Greger et al. 2017) on the basis that for example only two γ4 can co-assemble with AMPARs as seen with single-molecule photobleaching in live cells (Hastie et al. 2013).

In hippocampal CA1 neurons, almost exclusively expressing heteromeric AMPARs (Lu et al. 2009; Wenthold et al. 1996), 2-TARPs stoichiometry – together with 2 CNIHs – has been proven to be present (Gill et al. 2011). This stoichiometry of 2 TARPs and 2 CNIHs molecules assembling with CI-AMPARs might not be possible in CGCs due to the lack of cornichon homolog proteins (Schwenk et al. 2014). However, in CGCs, both γ2 and γ7 are expressed and thus γ7 could potentially be playing a role at somatic AMPARs. This is why we decided to explore resensitization in CGC somatic patches – a slow recovery of steady-state current after desensitization upon a prolonged agonist application firstly shown to be dependent on γ7 (Kato et al. 2007). Apart from γ7, also γ4 and γ8 show this exceptional resensitization signature (Gill et al. 2011) (but see (Carbone and Plested, 2016)). In the cerebellum the expression of γ4 and γ8 is negligible (Tomita et al. 2003) and CGCs do not express these TARPs (Fukaya et al. 2005). Besides, γ7 has been shown to selectively suppress CI-AMPARs in CGCs and its functional expression would be restricted to synaptic CP-AMPARs (Studniarczyk et al. 2013) although γ7 has been suggested not to have an important involvement in excitatory transmission in granule cells (Yamazaki et al. 2015). Our recordings from somatic patches of CGCs did not show the typical increase in steady-state current upon a 30 seconds in the presence of the full agonist AMPA, indicating that probably no functional γ7 molecules are present in somatic CI-AMPARs form CGCs in accordance with the poor role of this atypical TARP in granule neurons physiology shown to date (Studniarczyk et al. 2013; Yamazaki et al. 2015).

Another reason to explore resensitization in CGCs is that this property is a potential way to discriminate for a fully TARPed AMPAR since it has been suggested that resensitization is a fingerprint that indicates the presence of 4-TARPed AMPARs (Kato et al. 2010). Indeed, Purkinje cells from stargazer mice, with low γ7 stoichiometry lack this feature (Gill et al. 2011), indicating that saturation of TARPs is indispensable to present resensitization. The absence of resensitization in CGCs thus might be indicative of a 2-TARPed conformation. Despite recent reports showing new evidence that resensitization might not be a unique feature of TARPs γ4, γ7 and γ8 (Carbone and Plested, 2016), this does not exclude the possibility that resensitization is a characteristic signature of a fully TARPed receptor, thus reinforcing our conclusion of a 2-TARPed GluA2/GluA4c receptor in CGCs.

The results from different AMPAR-TARP combinations in recombinant systems showed that the 2-TARP-GluA2 condition displayed the slowest recovery from desensitization. AMPARs from somatic CGCs did show an even slower recovery rate compared with our expression systems data. We ruled out the possibility that the agonist used elicited a change in recovery kinetics (data not shown). One possibility of such slowdown in the recovery could be the presence of an “unknown” protein attached to the complex modulating the native AMPAR. In this regard, recovery from desensitization is strongly decreased either in neurons and heteromeric recombinant AMPARs when they are co-expressed with other auxiliary subunits of the CKAMP family as CKAMP39 (Farrow et al. 2015), CKAMP44 (von Engelhardt et al. 2010) or CKAMP52 (Klaassen et al. 20116) (*but see* (Farrow et al. 2015)). Interestingly, both CKAMP39 and 52 are highly expressed in the cerebellum (Farrow et al. 2015; Klaassen et al. 2016). Future work will elucidate whether apart from two γ2 molecules, CKAMPs play a role in AMPAR properties in CGCs.

A possible flaw in our study comes from the fact that in order to study AMPAR currents in CGCs, we have used the specific agonist AMPA to circumvent the stimulation of other glutamate-activated receptors – specially the kainate receptors present in CGCs (Bahn et al. 1994; Belcher and Howe, 1997). Certainly, the use of AMPA *vs.* glutamate may potentially alter the output values. When we checked for agonist variation on GluA4c with both agonists, neither weighted single-channel conductance nor open probability changed (data not shown). Previously, differences in channel conductance estimates have been described when using Kainate *vs.* AMPA to activate AMPARs (Jonas and Sakmann 1992; Swanson et al. 1997) but subconductance levels of CP-AMPARs have been shown to have similar values regardless of the use of AMPA or glutamate as agonist (Swanson et al. 1997). Furthermore, for GluA2/GluA4 similar conductance values have been described for both agonists 5.5-6 pS (Swanson et al. 1997), which match the ones observed in this and other studies (Jackson et al. 2011) using the agonist glutamate. Conversely, in our hands, AMPAR desensitization kinetics of GluA4c seemed to be significantly slower when AMPA was used as agonist (data not shown) despite the findings in previous reports that the kinetic properties of AMPA-activated GluA4 homomers were comparable to those activated by glutamate (Swanson et al. 1997). In principle, this might confuse the interpretation when comparing recordings in expression systems (glutamate used as agonist) with neurons (AMPA used as agonist). However, the kinetics of currents evoked with AMPA in CGCs were fast – indeed as rapid as the quicker responses observed in expression systems with glutamate. Therefore, it would be expected that the AMPAR-mediated responses in CGCs using the glutamate agonist would be faster than using AMPA as agonist– in any case kinetics would be overestimated – which still rules out that the possible stoichiometry present in CGCs were any of the slow combinations: 4 TARPs or 2 TARPs attached to GluA4c.

### Final remarks

AMPAR responses reckon on their subunit composition but also importantly rely on the type of auxiliary subunit/s accompanying them. This work adds important information on the extraordinary degree of functional variety in AMPARs by showing that AMPAR properties can be modulated differently depending on the number of TARPs, the type of AMPAR and the specific interactions that these auxiliary subunits set up with given GluAs. The broad number of possible combinations of pore-forming plus auxiliary subunits and stoichiometries that can be achieved on AMPARs generate a profuse diversity in glutamatergic responses in the brain. One of the major challenges scientists of the field will face is to resolve the exact composition of the AMPAR complex at different neurons.

## Methods

### Cell lines culture and transfection

tsA201 cells were used for electrophysiological experiments and were maintained and transfected with several constructs as described in (Gratacos-Batlle et al. 2014). More detailed information is available on Supplementary Information (Supplementary File 1).

### Cerebellar granule cells (CGCs) culture

Primary cultures of CGCs were prepared from pups on postnatal day 7–8 as previously described (Verdaguer et al. 2002). The cerebella from 8 to 10 mice pups were collected in 9.5 ml buffer containing 120 mM NaCl, 5 mM KCl, 25 mM HEPES, and 9.1 mM glucose. Thereafter, meninges were carefully removed and cerebella were dissected out, minced carefully with a blade, and dissociated at 37°C for 15 min with a solution containing 250 μg /ml trypsin. After 15 minutes, solution with 2.7 μg/ml DNAse and 8.32 μg/ml soybean trypsin Inhibitor (SBTI) was added. CGCs were separated from non-dissociated tissue by sedimentation and, finally, resuspended in basal medium Eagle’s (BME) supplemented with 10% inactivated fetal calf serum, 25 mM KCl, and gentamicin (5mg/ml), and plated onto poly-L-lysine-coated 24-well plates at a density of 300,000 cells/cm^2^. After 16–19h in culture, cytosine arabinoside was added to a final concentration of 10 μM to inhibit glial cell proliferation. Electrophysiological experiments were performed at 6 to 8 days after platting.

### Basic electrophysiological procedures

Recordings were performed from isolated transfected cells or cerebellar granule cells (CGCs) visualized with an inverted epifluorescence microscope (Axio-Vert.A1; Zeiss). Cells expressing EGFP and/or mCherry fluorescent proteins were selected for patch-clamp recordings. Macroscopic currents were recorded at room temperature (22-25°C) from outside-out membrane patches or from isolated whole cells using an Axopatch 200B amplifier and acquired using a Digidata 1440A interface board and pClamp 10 software (Molecular Devices Corporation, Sunnyvale, CA).

For all recordings the extracellular solution contained (in mM):145 NaCl, 2.5 KCl, 1 CaCl_2_, 1 MgCl_2_, 10 glucose and 10 HEPES (pH to 7.42 with NaOH). The extracellular control solution applied with the fast agonist application tool was composed by extracellular solution diluted 4% with miliQ H_2_O. The extracellular agonist solution applied with the fast agonist application tool was extracellular solution plus 2.5 mg/ml of sucrose with 10 mM glutamate for tsA201 recordings or 100 μM AMPA for tsA201 cells and CGC recordings. The intracellular pipette solution contained (in mM): 145 CsCl, 2.5 NaCl, 1 Cs-EGTA, 4 MgATP, and 10 HEPES (pH to 7.2 with CsOH). The polyamine spermine tetrahydrochloride (Sigma-Aldrich) was added to intracellular solution at 100 μM in all cases, which has been calculated to yield free concentrations in the physiological range (Soto et al. 2007).

Patch pipettes were fabricated from borosilicate glass (1.5 mm o.d. and 0.86 mm i.d.; Harvard Apparatus, Edenbridge, UK) by using a Horizontal puller (Sutter P-97) with several resistances depending on the configuration used.

Recordings were performed in the whole-cell or outside-out configurations, where different parameters as recovery from desensitization, desensitization kinetics, single channel conductance, peak open probability, intracellular polyamine block or percentage of block by non-competitive antagonists was tested as explained in detail in Supplementary File 1.

### Statistical Analysis

Analysis of current waveforms and curve fitting was performed with IGOR Pro 6.06 (Wavemetrics) using NeuroMatic 2.03 ((Rothman and Silver, 2018); http://www.neuromatic.thinkrandom.com). Statistical analysis was performed using GraphPad Prism version 8.0.1 for Mac OS X (GraphPad Software, San Diego California USA, www.graphpad.com). Comparisons between two groups were performed using the parametric Student’ *t*-test for data following a normal distribution or using the non-parametric Mann-Whitney U test for comparisons between groups in which one of them did not follow a normal distribution. Differences were considered significant as follows: *p < 0.05, **p<0.01, ***p<0.001 and ****p<0.0001. Extended methodology is available in Supplementary File 1 (Animals and housing, constructs, electrophysiological configurations used and analysis performed).

## Acknowledgements

We would like to thank Francesc Sureda (Universitat Rovira i Virgili) for technical assistance with cerebellar granule cell cultures and Jon Giblin for English revision.

## SOURCE DATA FILES FOR FIGURES

Figure 1 - Source Data 1

Figure 2 - Source Data 2

Figure 3 - Source Data 3

Figure 4 - Source Data 4

Figure 5 - Source Data 5

Figure 6 - Source Data 6

Figure 7 - Source Data 7

Figure S1 - Source Data S1

Figure S2 - Source Data S2

## SUPLEMMENTARY FILES

Supplementary File 1

---

**Figure S1:**
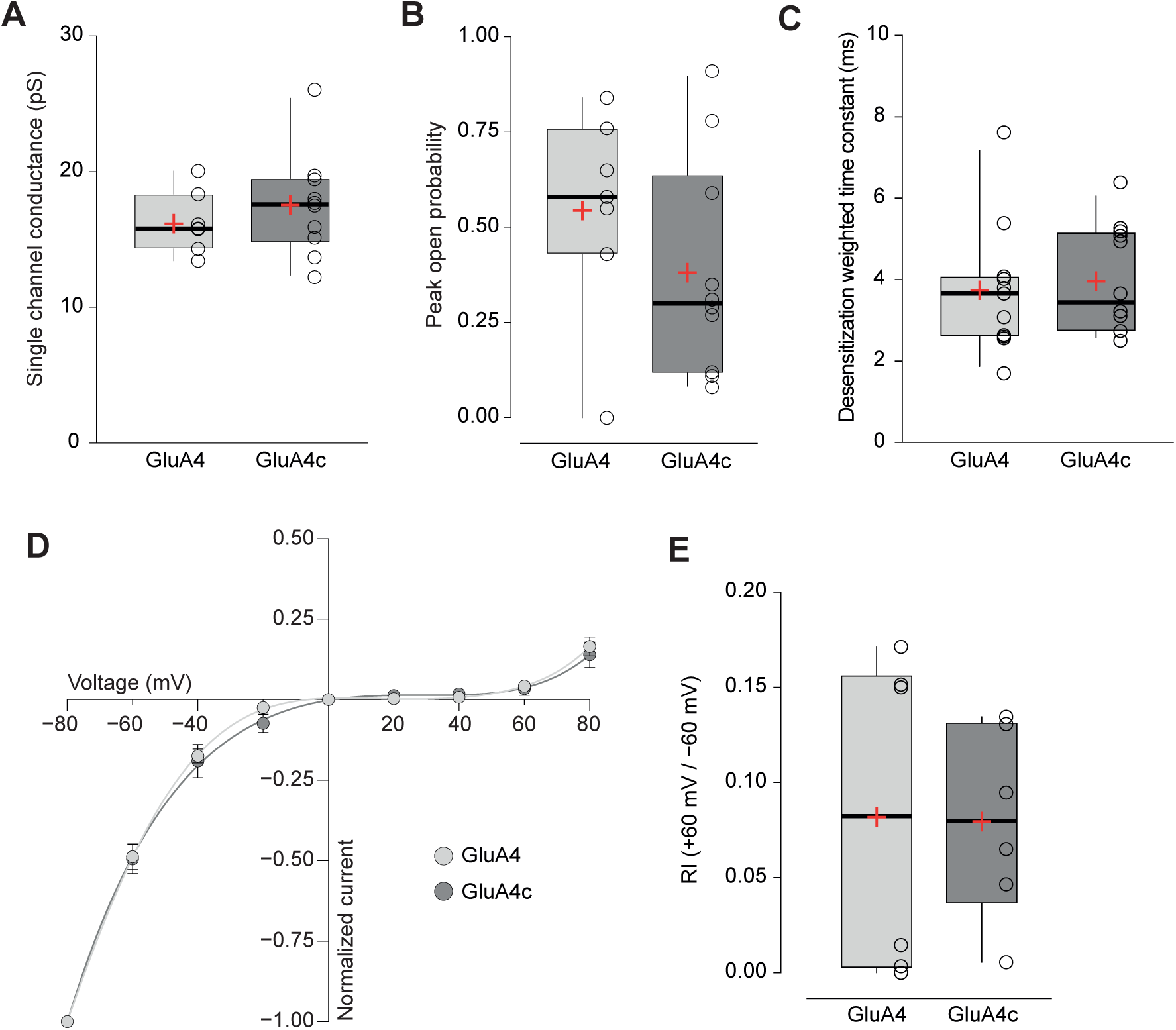
Short and long isoforms of GluA4 show the same electrophysiological behaviour. **A**. Pooled data showing single channel conductance from GluA4 (light grey) and GluA4c (dark grey) homotetramers. Box-and-whiskers plots represent percentiles, median and average as stated in Figure 1c. **B**. Data showing peak open probability from GluA4 and GluA4c tetramers using the same colour pattern as in figure 5a. **C**. Weighted time constant of desensitization (τw, des) from GluA4 and GluA4c tetramers. **D**. Inwardly rectifying I-V relationships constructed from peak currents evoked by glutamate (100 ms, 10 mM) applied to outside-out patches from tsA201 cells containing GluA4 (light circle) or GluA4c (dark circle). Fittings are adjusted to a 6th order polynomial function. **E**. RI (+60mV/-60mV) values from GluA4 and GluA4c tetramers. The data from this figure containing statistical tests applied, exact sample number, p values and details of replicates are available in “Figure S1 - Source Data S1”.

**Figure S2:**
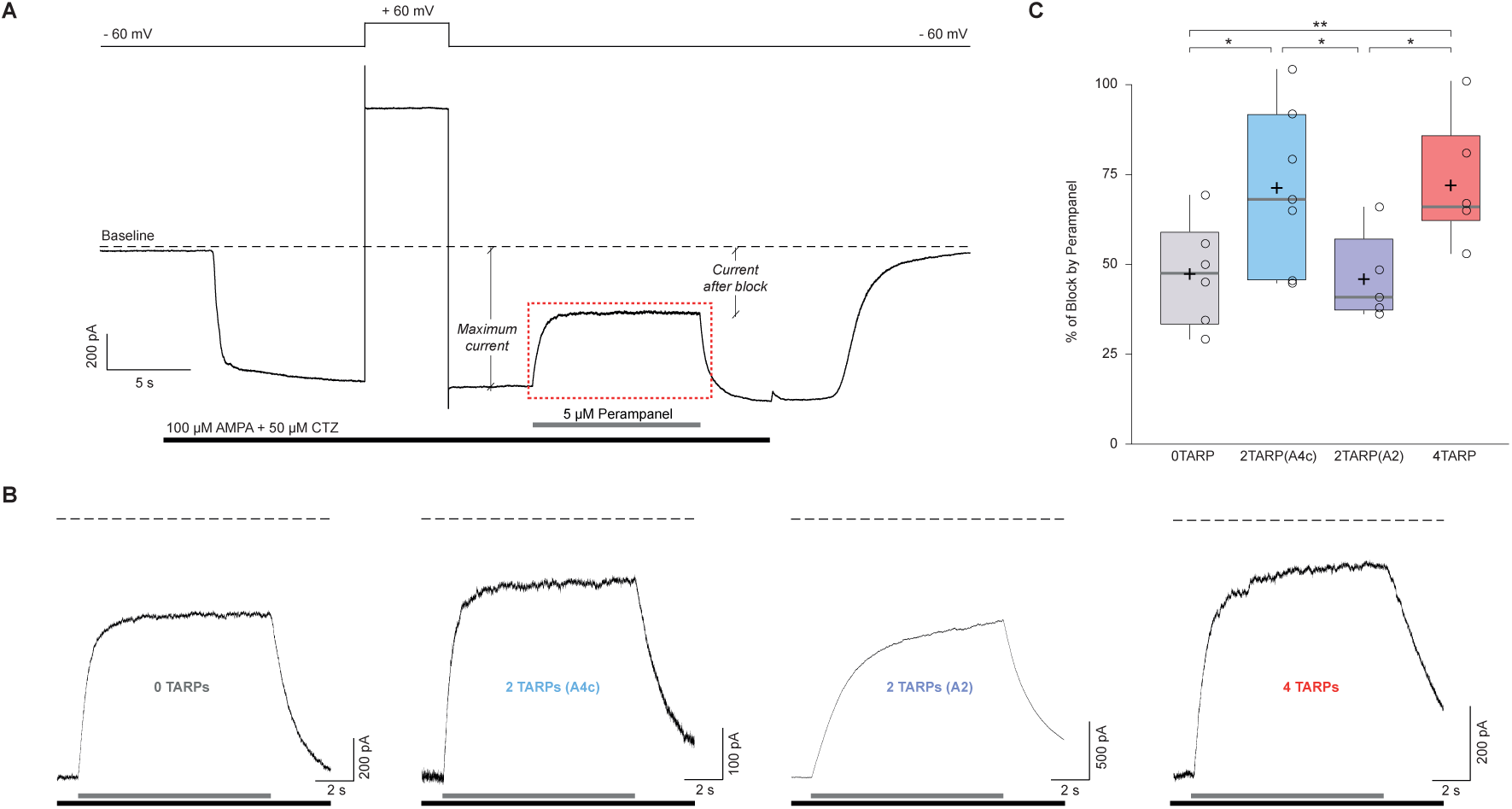
The block of the non-competitive antagonist perampanel varies with AMPAR-TARP stoichiometry. **A**. Representative whole-cell recording showing the protocol used to determine the percentage of perampanel block. This trace corresponds to a cell expressing GluA2 and GluA4c. Cells were clamped at −60 mV during the recording with a 5 second pulse duration at +60 mV to asses for GluA2 presence. Black bar shows application time of AMPA 100 μM + Cyclothiazide 50 μM to tsA201 cells expressing different AMPAR-TARP combinations. Grey bar shows rapid application (< 1ms) of Perampanel at 5μM. Baseline is pointed with dashed line. Maximum current and current after block are displayed. The dashed red frame represents the magnification part shown in panel b. **B**. Representatives traces of Perampanel blocking from each condition following the same colour pattern as in Figure 5. Under each trace the black bar shows AMPA + CTZ and grey bar shows application of Perampanel. **C**. Pooled data for the percentage of blocking by Perampanel. Box-and-whiskers plots as in Figure 1c. The data from this figure containing statistical tests applied, exact sample number, p values and details of replicates are available in “Figure S2 - Source Data S2”.

## Supplementary Material and Methods

### 1.1 Animals and housing

C57BL/6N wild-type mice were housed in cages with free access to food and water and were maintained under controlled day–night cycles in accordance with the NIH Guide for the Care and Use of Laboratory Animals, the European Union Directive (2010/63/EU), and the Spanish regulations on the protection of animals used for research, following a protocol approved and supervised by the CEEA-UB (Ethical Committee for Animal Research) from University of Barcelona with the license number OB117/16, of which DS is the responsible researcher.

### 1.2 Cell lines culture and transfection

In this study, we have used tsA201 cells, a cell line derived from HEK293 cells stably transfected with the temperature sensitive gene for SV40 T-antigen to allow plasmid replication using the SV40 origin and hence to produce high levels of recombinant proteins (Sigma catalog #85120602). Cells were maintained as described in Gratacòs-Batlle et al., 2015 ^29^.

Cells were plated into poly-D-lysine coated coverslips 24 h before transfection at a density of 1.5 × 10^6^ cells/coverslips. Cells were then transiently transfected with 0.8-1 μg total cDNA using PEI reagent (1 mg/ml) in a 3:1 ratio (PEI:DNA) according to the manufacturer’s directions. The ratio of cDNA used in each condition varied depending on the set of experiments.

### 1.3 Constructs

GluA1, GluA2 and GluA4 cDNAs (rat, flip isoforms) were old gifts from S. Heinemann (Salk Institute, La Jolla, CA, USA) and P. Seeburg (Max Planck Institute, Heidelberg, Germany).

For this work we used the short version of GluA4, namely GluA4c – first described in 1992 ^30^ – in its flip form. The GluA4c subunit was cloned from mRNA obtained from adult rat cerebellum (*Rattus norvegicus*) cerebellum into a pIRES-mCherry plasmidic vector. The primers used were the following:

Primer Forward (5’-3’): GCGC GCT AGC ATG AGG ATT TGC AGG CAG ATT (GCGC GCT AGC restriction site cloned using NheI-HF enzyme, catalog: NEB #R3131S).

Primer Reverse (5’-3’): CGCGG CTC GAG ATT CTT AAT ACT TTC GGT TCC A (CGCGG CTC GAG restriction site cloned using XhoI-HF enzyme, catalog: NEB #R0146S).

GluA1:γ2 and GluA2:γ2 tandem proteins (into pIRES-GFP vectors) were a generous gift from Ian Coombs (UCL, London, UK). GluA4c:γ2 tandem was subcloned into a pIRES-mCherry vector from the GluA4c and the γ2 plasmidic vectors. All constructs have been fully sequenced.

### 1.4 Electrophysiology

#### 1.4.1 Whole-cell recordings

Whole-cell recordings were made from isolated cells using electrodes with a resistance of 3–5MΩ, giving a final series resistance of 5–15 MΩ. Voltage was held at −60 mV unless otherwise stated. Currents were low-pass filtered at 2 kHz and digitized at 5 kHz. Receptors were activated by bath application of 10 mM glutamate plus 50 μM cyclothiazide (CTZ) to prevent receptor desensitization.

#### 1.4.2 Fast agonist application into outside-out patches

Outside-out patches were obtained using electrodes with a resistance of 5–10 MΩ. Rapid solution switching at the patch was carried out by piezoelectric translation of a theta-barrel application tool made from borosilicate glass (1.5mm o.d.; Sutter Instruments) mounted on a piezoelectric translator (P-601.30; Physik Instrumente). Control and agonist solutions flowed continuously through the two barrels of the theta glass and solution exchange occurred when movement of the translator was triggered by a voltage step (pClamp). To enable visualization of the solution interface and to allow measurement of the solution exchange 2.5 mg/ml sucrose was added to the agonist solution and the control solution was diluted by 4%. Currents were activated by 10mM glutamate and were low-pass filtered at 10 kHz and digitized at 50 kHz. At the end of each experiment, the adequacy of the solution exchange was assessed by destroying the patch and measuring liquid-junction current at the open pipette; the 10–90% rise time was always <400 μs.

#### 1.4.3 Recovery from desensitization

To study AMPAR recovery from desensitization, a two-pulse protocol (25 ms each) was used in which a first pulse was applied followed by a second pulse at different time intervals (from 25 ms to 625 ms). The paired pulses were separated 1s to allow full recovery from desensitization. To estimate the percentage of recovery, the magnitude of peak current at the second pulse (P2) was compared with the first one (P1).

#### 1.4.4 Non-stationary fluctuation analysis (NSFA)

To infer channel properties from macroscopic responses, glutamate (10 mM) was applied onto outside-out patches (100 ms duration at 1 Hz) and the ensemble variance of all successive pairs of current responses was calculated using IGOR Pro 6.06 (Wavemetrics, OR) and NeuroMatic ^33^. The single-channel current (*i*), total number of channels (N) and maximum open probability (P_o,max_) were then determined by plotting this ensemble variance (*σ*^2^) against mean current (I) and fitting with a parabolic function:

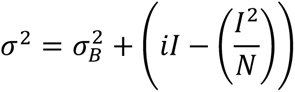

where 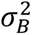 is the background variance. Along with the expected peak-to-peak variation in the currents due to stochastic channel gating, some responses showed a gradual decrease in peak amplitude (*run-down* of the current). The mean response was calculated from epochs containing 50 to 350 stable responses, which were identified by using a Spearman rank-order correlation test (NeuroMatic). The weighted-mean single-channel conductance was calculated from the single-channel current and the holding potential (being −0.055 V after corrected for liquid-junction potential). P_o,max_ was estimated by dividing the average peak current by theoretical maximum current (*i*N).

#### 1.4.5 Kinetics of desensitization

The kinetics of desensitization of glutamate-evoked responses and the kinetics of recovery from desensitization were fitted according to a double-exponential function to calculate the weighted time constant (*τ*_*w,des*_):

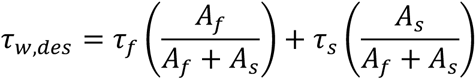

where *A*_*f*_ and *τ*_*s*_ are the amplitude and time constant of the fast component of recovery and *A*_*s*_ and *τ*_*s*_ are the amplitude and time constant of the slow component.

#### 1.4.6 Current-voltage relationships

In order to study the degree of spermine block of CP-AMPARs at different membrane potentials we applied 10 mM glutamate onto outside-out membrane patches at different holding potentials (from −80 mV to +80 mV in 20 mV increments) and the peak current was used to construct the current-voltage relationship.

The rectification index (*RI*) was defined as the absolute value of glutamate-evoked current at +60 mV divided by that at −60 mV:

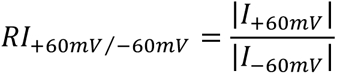

